# Rab40c Regulates Focal Adhesions and Protein Phosphatase 6 Activity by Controlling ANKRD28 Ubiquitylation and Degradation

**DOI:** 10.1101/2021.12.06.471409

**Authors:** Ke-Jun Han, Rytis Prekeris

## Abstract

Rab40c is a SOCS box–containing protein which binds Cullin5 to form a ubiquitin E3 ligase complex (Rab40c/CRL5) to regulate protein ubiquitylation. However, the exact functions of Rab40c remain to be determined, and what proteins are the targets of Rab40c-Cullin5 mediated ubiquitylation in mammalian cells are unknown. Here we showed that in migrating MDA-MB-231 cells Rab40c regulates focal adhesion’s number, size, and distribution. Mechanistically, we found that Rab40c binds the protein phosphatase 6 (PP6) complex and ubiquitylates one of its subunits, ankyrin repeat domain 28 (ANKRD28), thus, leading to its lysosomal degradation. Furthermore, we identified that phosphorylation of FAK and MOB1 is decreased in Rab40c knock-out cells, which may contribute to focal adhesion site regulation by Rab40c. Thus, we propose a model where Rab40c/CRL5 regulates ANKRD28 ubiquitylation and degradation, leading to a decrease in PP6 activity, which ultimately affects FAK and Hippo pathway signaling to alter focal adhesion dynamics.

## INTRODUCTION

Cell migration is a fundamental cellular function that is involved in many important biological processes, including embryological development, tissue formation, wound healing, and cancer metastasis. In response to extracellular and intracellular signals, migratory cells reorganize their cytoskeleton and endocytic transport to form actin-dependent migratory cell protrusions, such as lamellipodia, and establish a front-to-rear polarity (Maritzen, Schachtner et al., 2015, Seetharaman & Etienne-Manneville, 2020, SenGupta, Parent et al., 2021, Shellard & Mayor, 2020). By interaction with the extracellular matrix (ECM), cells can form integrin-based macromolecular adhesive structures called focal adhesions (FAs) (Burridge, 2017, Burridge & Guilluy, 2016, Legerstee & Houtsmuller, 2021, Mishra & Manavathi, 2021). FAs include numerous scaffoldings and signaling proteins, such as talin, vinculin, zyxin, paxillin, p130Cas, and α-actinin, that regulate FA formation and disassembly during cell migration (Ibata & Terentjev, 2020, Llanses Martinez & Rainero, 2019, Stutchbury, Atherton et al., 2017). One of the main functions of FAs is to physically connect the cellular actin cytoskeleton to ECM, therefore sensing, integrating, and transducing extracellular signaling. In addition, FAs can serve as anchor points to generate tensional forces to push cells forward during cell migration (Burridge & Guilluy, 2016, Parsons, Horwitz et al., 2010, Seetharaman & Etienne-Manneville, 2020, Yamada, Collins et al., 2019).

FAs formation is initiated by ECM binding to cell surface receptors, primarily integrins, at the leading edge (Legerstee & Houtsmuller, 2021, Schumacher, Vazquez Nunez et al., 2021). Newly formed nascent adhesions gradually grow and change their protein composition to mature into FAs. FAs usually localize at the cell periphery, where they associate with the ends of stress fibers (Burridge & Guilluy, 2016, Zaidel-Bar, Cohen et al., 2004). With nascent adhesion formation at the leading edge, the FAs at the cell rear need to be disassembled to promote rear end retraction and efficient cell migration. Therefore, FAs are highly dynamic, and their number, size, and distribution can rapidly change in response to internal or external signals. It is well established that the dynamic process of FAs is under the regulation of protein tyrosine kinases such as focal adhesion kinase (FAK) and small GTPases of the Rho family (Katoh, 2017, Nobes & Hall, 1999, Tapial Martinez, Lopez Navajas et al., 2020). Recently, Hippo signaling pathways also have been suggested to regulate cell migration by controlling FA dynamics and mediating mechano-sensing of changes in ECM stiffness and composition (Nardone, Oliver-De La Cruz et al., 2017, Rausch & Hansen, 2020, Wang, Englund et al., 2021)). Although extensively studied, our current understanding of the molecular mechanisms underlying FA dynamics is still limited.

Rab GTPases are key regulators of membrane trafficking and play an important role in cell migration. We previously demonstrated that the Rab40 subfamily of small GTPases is required for breast cancer cell invasion by promoting ECM degradation and invadopodia formation (Jacob, Jing et al., 2013, Jacob, Linklater et al., 2016, Linklater, Duncan et al., 2021), although it remains to be fully understood how proteins within the Rab40 family function and what molecular machinery is governing Rab40-dependent cell migration and invasion. Rab40 is a unique subfamily of small monomeric GTPases that include four closely related proteins: Rab40a, Rab40al, Rab40b, and Rab40c, and is characterized by the presence of suppressors for the cytokine signaling (SOCS) box at their C-terminal (Homma, Hiragi et al., 2021, Klopper, Kienle et al., 2012, Pereira-Leal & Seabra, 2000). The SOCS box in other proteins has been shown to bind Cullin5–ElonginB/C to form ubiquitin E3 ligase complex (CRL5), used to mediate target protein ubiquitination and degradation (Kile, Schulman et al., 2002, Linossi & Nicholson, 2012), and we recently demonstrated that Rab40b binds CRL5 to ubiquitylate EPLIN and promote cell migration by altering focal adhesion and stress fiber formation (Linklater et al., 2021). Furthermore, Rab40a was implicated in mediating proteasomal degradation of RhoU, thus, regulating FA dynamics (Dart, Box et al., 2015). All these findings suggest that the Rab40 subfamily of small monomeric GTPases may have evolved to regulate actin dynamics and FA turn-over by mediating ubiquitylation and degradation of a specific subset of proteins that regulate cell migration.

In this study we focus on investigating the functions of Rab40c, since it remains to be determined whether Rab40c regulates FA dynamics. Additionally, it remains unclear what are the targets of Rab40c-dependent ubiquitylation and degradation. Here, we show that Rab40c regulates the size, location, and number of FAs in breast cancer cells, whilst also demonstrating that Rab40c interacts with ankyrin repeat domain 28 protein (ANKRD28), which is a scaffolding subunit of heterotrimeric protein phosphatase PP6 complex. Importantly, ANKRD28-containing PP6 complex inhibits FA formation, and Rab40c directly regulates ubiquitination and degradation of ANKRD28 in breast cancer cells. Finally, we found that Rab40c regulates the Hippo signaling pathway, possibly through MOB1 dephosphorylation by an ANKRD28-containing PP6 subcomplex. Based on all this data, we propose that Rab40c/CRL5 contributes to regulation of FAs by inhibiting the formation and activity of ANKRD28-containing PP6 subcomplexes, which in turn regulates Hippo signaling and mechanosensing.

## RESULTS

### Rab40c regulates focal adhesion formation

Rab40b small monomeric GTPases have recently emerged as regulators of localized ubiquitylation of several proteins involved in mediating cell migration (Linklater et al., 2021). Rab40c is another member of the Rab40 sub-family of proteins that is defined by the presence of SOCS box and their ability to mediate Cullin5-dependent protein ubiquitylation. However, despite its close similarity to Rab40b (Duncan, Lencer et al., 2021), it remains unclear whether Rab40c plays any role in regulating cell migration, and whether Rab40c and Rab40b have overlapping functions. Since Rab GTPases are key regulators of intracellular membrane trafficking and localization of different membrane compartments, we first decided to examine the subcellular localization of human Rab40c. To that end, we created MDA-MB-231 cell line stably expressing GFP-tagged human Rab40c, and analyzed the distribution of GFP-Rab40c by immunofluorescence microscopy. As shown in Figure1A, while the majority of GFP-Rab40c is present in the cytosol, a subpopulation of GFP-Rab40c can clearly be observed at the front edge of the lamellipodia where it colocalizes with actin ruffles. GFP-Rab40c also accumulates and colocalizes with GM130 and VAMP4 (Supplemental Fig. 1), two Golgi markers, suggesting that in addition to potentially regulating lamellipodia dynamics, GFP-Rab40c may also regulate protein transport from Golgi to the plasma membrane.

**Figure 1.**
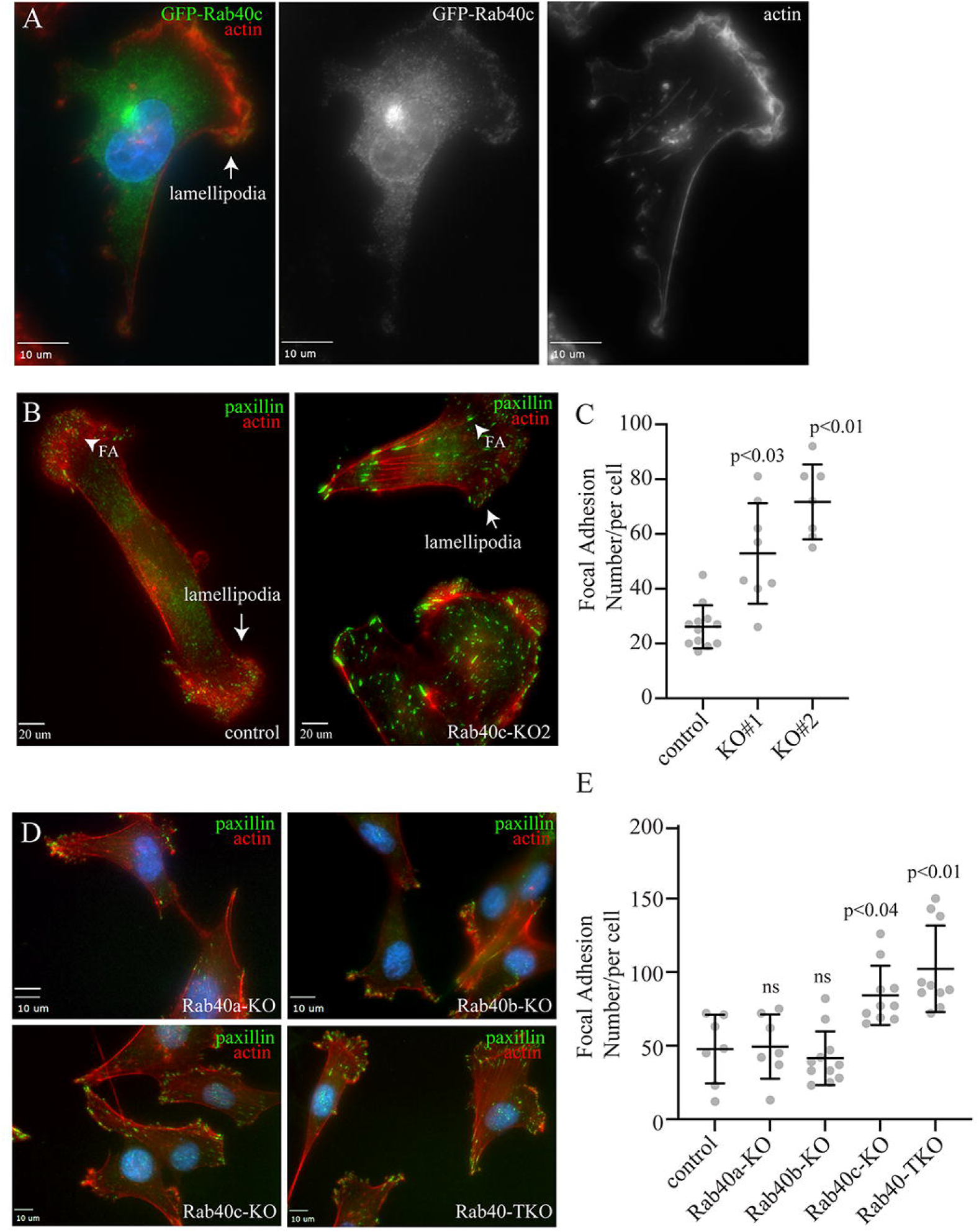
Rab40c regulates focal adhesions. (A) MDA-MB-231 cells stably expressing GFP-Rab40c were plated on collagen-coated coverslips and then fixed and stained with phalloidin-AlexaFluor594. Arrow points to the lamellipodia. (B) Control or Rab40c-KO MDA-MB-231 cells fixed and stained with phalloidin-AlexaFluor594 (red) and and anti-paxillin antibodies (green). Arrows point to the lamellipodia, and arrowheads point to FAs. (C) Quantification of number of FAs per cell for control and Rab40c KO cells. n ≥ 10 cells per condition. Data shown are means and standard deviations derived from two independent experiments. (D) Control, Rab40a-KO, Rab40b-KO, Rab40c-KO and Rab40a/b/c-KO (Rab40-TKO) MDA-MB-231 cells were fixed and stained with phalloidin-AlexaFluor594 (red) and anti-paxillin antibodies (green). (E) Quantification of number of FAs per cell for control, Rab40a-KO, Rab40b-KO, Rab40c-KO and Rab40-TKO cells. n ≥ 10 cells per condition. Data shown are means and standard deviations derived from two independent experiments.

Our previous study showed that knock-out of all three Rab40 isoforms (Rab40a, Rab40b, and Rab40c) inhibited cell migration in part by increasing the number and size of focal adhesion sites (FAs) (Linklater et al., 2021). What remains unclear is which Rab40 sub-family members may be involved in directly regulating FA dynamics. To test whether Rab40c has effects on FAs, we generated Rab40c knockout cell lines by CRISPR/Cas-mediated genome editing, which have been validated by genomic sequencing and western blot (Supplemental Fig. 2B-C). These cells were then stained with an anti-paxillin antibody to assess the steady-state number and distribution of FAs. As previously reported in the control cells, paxillin positive dot-like FAs were mostly present at the cell periphery, especially leading-edge lamellipodia (Fig. 1C). In contrast, within Rab40c-depleted cells, an increased quantity with larger paxillin-positive FAs were observed (Fig. 1C). Importantly, FAs were not limited to the periphery of the cell, but instead were scattered throughout the plasma membrane (Fig. 1C). Quantification of the number of FAs per cell shows a significant increase of FAs in Rab40c KO cells (Fig.1D). This change seems to be mediated predominately by Rab40c-KO because neither Rab40a nor Rab40b KOs alone led to an increase in FA number and size. Consistent with this and with our previously published data (Linklater et al., 2021), knocking-out all three Rab40 isoforms (Rab40a, Rab40b, and Rab40c; Rab40-TKO) did not further increase the number of FAs as compared to Rab40c-KO (Fig.1E-F).

**Figure 2.**
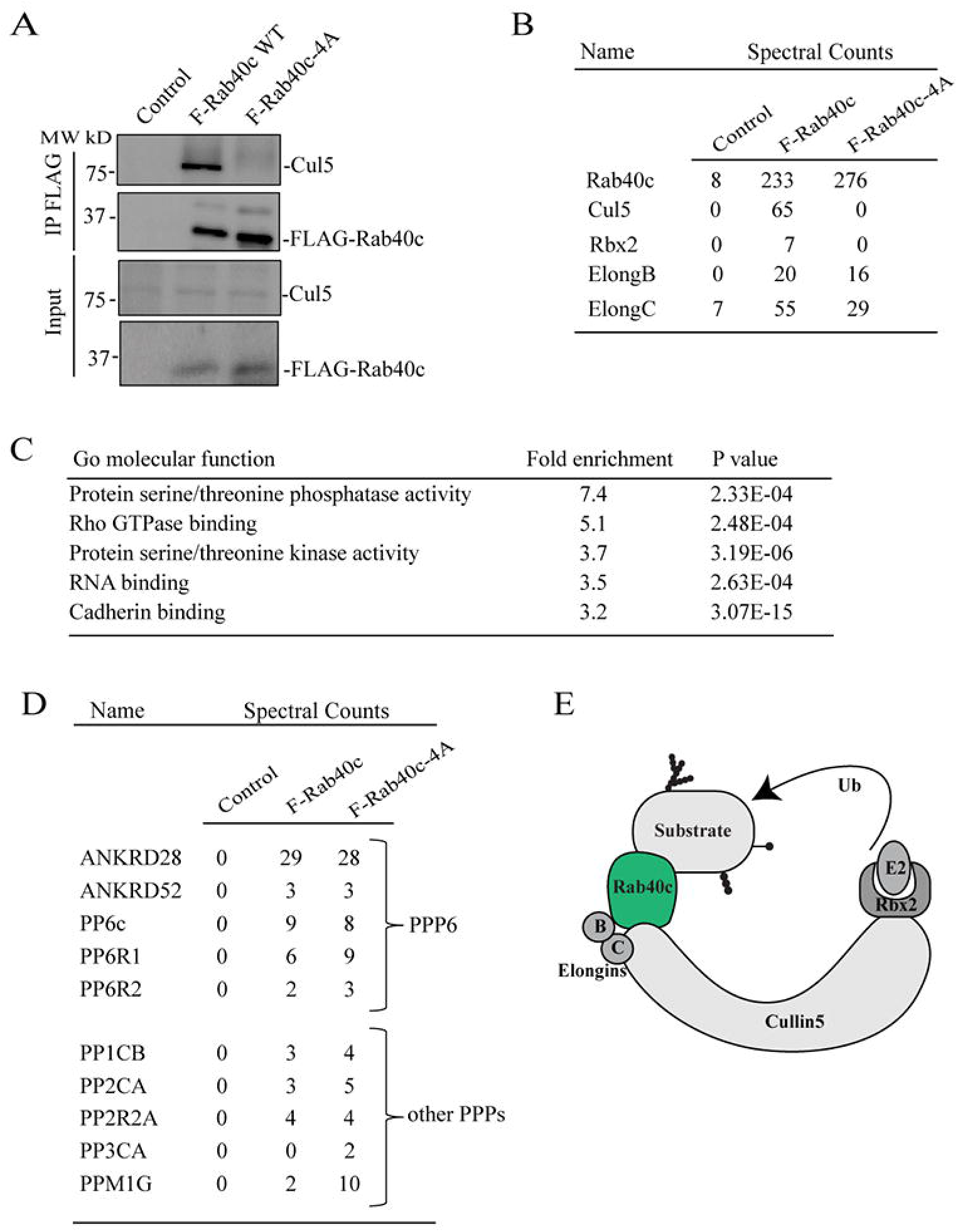
Identification of Rab40c-interacting proteins. (A) Cell lysates from control, FLAG-Rab40c and FLAG-Rab40c-4A expressing cells were immunoprecipitated with anti-FLAG antibody. Immunoprecipitates and cell lysates then were then blotted with anti-FLAG or anti-Cul5 antibodies. (B) The list of Cul5 ligase complex (CRL5) subunits identified by mass spectrometry from FLAG-Rab40c and FLAG-Rab40c-4A expressing cells. (C) Gene Ontology (GO) enrichment analysis of proteins identified by mass spectrometry from FLAG-Rab40c and FLAG-Rab40c-4A immunoprecipitates. (D) List of phosphoprotein phosphatases identified by mass spectrometry from FLAG-Rab40c and FLAG-Rab40c-4A immunoprecipitates. (E) A model showing Rab40c-CRL5 E3 ligase complex. Ub-ubiquitin.

### Identification of Rab40-interacting proteins

To understand how Rab40c contributes to regulation of FAs, we next sought to identify Rab40c-interacting partners. Rab40c has a SOCS box at its C-terminus, and it is well established that a highly conserved LPLP motif in the SOCS box is necessary for the binding to Cullin5 (Cul5). To examine whether this motif in Rab40c is also important for the binding to Cul5, we mutated the LPLP sequence to AAAA (FLAG-Rab40c-4A) and established MDA-MB-231 cell lines stably expressing either FLAG-Rab40c or FLAG-Rab40c-4A. Then, FLAG tagged proteins were immunoprecipitated with an anti-FLAG antibody. As shown in Figure 2A, endogenous Cul5 co-precipitated with FLAG-Rab40c. However, FLAG-Rab40c-4A has lost its ability to bind Cul5, confirming that the LPLP motif mediates Rab40c binding to Cul5.

To identify proteins that bind to Rab40c, we co-immunoprecipitated FLAG-Rab40c with either anti-FLAG antibody–conjugated beads or control IgG beads, followed by analysis using mass spectrometry. As expected, Cul5 and Rbx2 both co-immunoprecipitated with FLAG-Rab40c, but not IgG control or FLAG-Rab40c-4A (Fig. 2B). Furthermore, Elongin B/C, two known components of CRL5 ubiquitin E3 ligase complex, were also identified in the elutes from both FLAG-Rab40c and FLAG-Rab40c-4A, consistent with previous work that Elongin B/C binding to Rab40c is independent of LPLP motif and Cul5 (Fig. 2B). All other putative Rab40c-binding proteins identified by mass spectrometry analysis were then filtered to eliminate possible contaminants. Only candidates that were absent in IgG control, but present in both FLAG-Rab40c and FLAG-Rab40c-4A, were identified as putative Rab40c interactors. Furthermore, all RNA, DNA, and mitochondria-binding proteins were eliminated as putative contaminants (Supplemental Table 1).

Gene Ontology (GO) enrichment analysis of putative Rab40c interactors reveals strong enrichment for protein serine/threonine phosphatases and serine/threonine kinases, among the putative Rab40c-binding proteins (Fig. 2C), suggesting that Rab40c may regulate FA-dependent signaling. Specifically, Ankyrin repeat domain 28 (ANKRD28), a subunit of protein phosphatase 6 (PP6), was one of the highly enriched proteins. Importantly, other components of PP6, including ANKRD52, catalytic PP6 subunit (PP6c), and PP6R1/2, were all identified in the Rab40c precipitates (Fig. 2D), indicating that the PP6 complex may interact and be regulated by Rab40c. Other PPs such as PP1CB, PP2CA, and PPM1G were also identified as putative Rab40c-binding proteins (Fig. 2D), however, in the rest of this study we will focus on interaction between PP6 complex and Rab40c.

### Rab40 interacts with the PP6 complex that contains ANKRD28 and PP6R1 subunits

The PP6 holoenzyme is a hetero-trimeric complex formed by the catalytic subunit PP6c, one of the regulatory subunits of PP6R1, 2, or 3, and one of an ankyrin repeat-domain containing protein ANKRD28, 44, or 52 (Fig. 3A). PP6Rs and ANKRDs are generally considered to be regulatory and scaffolding subunits that determine the localization and specificity of PP6 complexes (Nilsson, 2019, Ohama, 2019). Thus, it is now widely accepted that there are several PP6 complexes, and that the composition of these complex is what defines PP6 function and specificity for substrate proteins. To confirm that Rab40c binds to the PP6 complex, we over-expressed FLAG-Rab40c and FLAG-Rab40c-4A in 293T cells, followed by precipitation with anti-FLAG antibodies, and immunoblotting for endogenous ANKRD28 and PP6R1-3, two PP6 regulatory/scaffolding subunits that were most abundant in FLAG-Rab40c proteomic analysis (Fig. 2D). Consistent with our proteomics data, we found that ANKRD28 and PP6R1 both interacted with FLAG-Rab40c and FLAG-Rab40c-4A (Fig. 3B). To further confirm Rab40c and PP6 complex interaction, we immunoprecipitated endogenous Rab40c from 293T cells and found that ANKRDs and PP6R1, but not PP6R2 and PP6R3, were pulled out by an anti-Rab40c antibody (Fig. 3C; Supplemental Fig. 2A).

**Figure 3.**
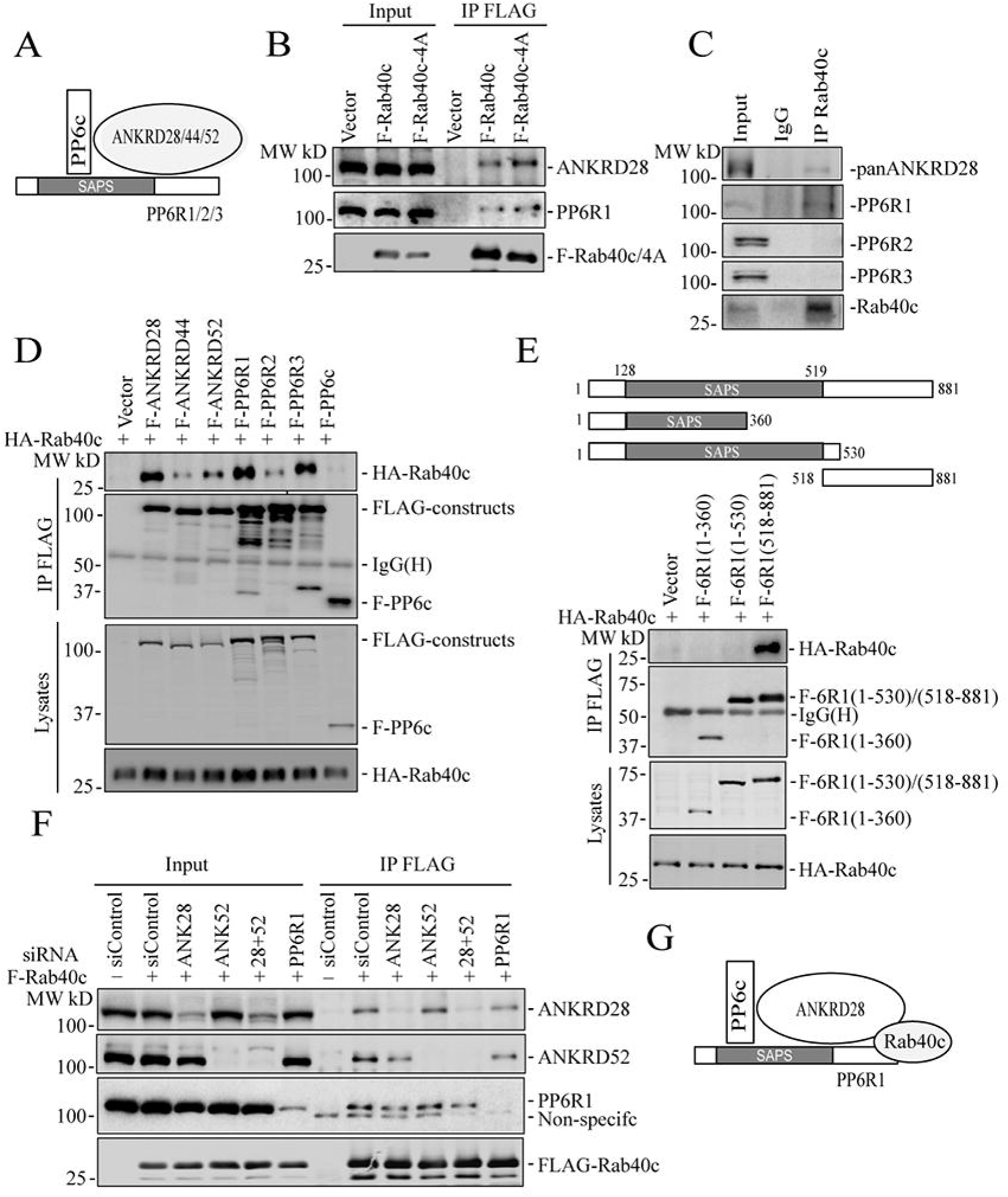
Rab40c interacts with the PP6 complex. (A) A model showing the PP6 complex. (B) 293T cells were transfected with control (empty plasmid), FLAG-Rab40c or FLAG-Rab40c-4A plasmids and then cell lysates were immunoprecipitated with an anti-FLAG antibody. Cell lysates and immunoprecipitates were then immunoblotted with anti-ANKRD28, anti-PP6R1 or anti-FLAG antibodies. (C) 293T cell lysates were immunoprecipitated with anti-Rab40c and immunoprecipitate was then blotted with anti-Rab40c, anti-panANKRD, anti-PP6R1, anti-PP6R2, or anti-PP6R3 antibodies. (D) 293T cells were co-transfected with HA-Rab40c and control (empty plasmid) or one of FLAG-tagged PP6 components. Cell lysates were then immunoprecipitated with an anti-FLAG antibody. Cell lysates and precipitates were immunoblotted with anti-HA or anti-FLAG antibodies. (E) Top panels: a schematic diagram of PP6R1 deletion mutants. Lower panels: 293T cells were co-transfected with HA-Rab40c and control or one of FLAG-tagged PP6R1 deletion mutants. Cell lysates were then immunoprecipitated with an anti-FLAG antibody. Cell lysates and precipitates were immunoblotted with anti-HA or anti-FLAG antibodies. (F) 293T cells were co-transfected with indicated siRNA(s) and FLAG-Rab40c. Cell lysates were then immunoprecipitated with an anti-FLAG antibody. Cell lysates and immunoprecipitates were immunoblotted with indicated antibodies. (G) A model showing proposing that Rab40c interacts with PP6R1 and ANKRD28 subunits of PP6 complex.

To identify what PP6 subunits mediate interaction with Rab40c, we overexpressed HA-Rab40c with all seven FLAG-tagged PP6 subunits in 293T cells and performed individual immunoprecipitations using an anti-FLAG antibody. Although we precipitated comparable amounts of various PP6 subunits, and HA-Rab40c was detected in all of immunoprecipitations to varying degrees, the highest amounts of HA-Rab40c co-precipitated with ANKRD28, PP6R1, and PP6R3 (Fig. 3D). Interestingly, PP6R3 was also identified in Rab40c proteomic analysis (Supplemental Table 1) but was eliminated from further analysis since some of the PP6R3 was also detected in IgG control. Taken together, this data suggests that Rab40c preferentially interacts with PP6 complexes containing PP6R1/ANKRD28 subunits.

It was proposed that PP6R1 functions as a scaffolding protein for PP6 holoenzyme assembly (Guergnon, Derewenda et al., 2009, Stefansson, Ohama et al., 2008), therefore, we set out to determine which region of PP6R1 binds to Rab40c. To that end, we generated a series of FLAG-tagged PP6R1 deletion mutants including PP6R1(1-360), PP6R1(1-530), and PP6R1(518-881) (Fig. 3E), and then individually co-transfected all these constructs with HA-Rab40c, followed by co-IP with anti-FLAG and blotting with anti-HA antibodies. The results suggest that the C-terminal region of PP6R1 spanning amino acids 518–881 mediates its interaction with Rab40c (Fig. 3E). Importantly, PP6R1(518-881) is outside of the SAPS domain that mediates PP6c binding to the PP6R1, suggesting that Rab40c may be able to interact with the entire PP6 holoenzyme rather than just PP6R1.

PP6 complex contains both PP6R1 and ANKRD28 subunits, and any one of them can recruit Rab40c to the PP6c/ANRD28/PP6R1 complex (Fig. 3A). To identify which subunits are required for PP6 complex binding to Rab40c, we transfected 293T cells with FLAG-Rb40c and scrambled siRNA, or siRNAs targeting ANKRD28, ANKRD52, and PP6R1. FLAG-Rab40c was then immunoprecipitated, and the immunoprecipitates were analyzed for the presence of ANKD28, ANKRD52, or PP6R1 by Western blotting. As shown in (Fig. 3F), siRNA-mediated knockdown effectively reduced individual protein levels > 80%. However, none of these knock-downs completely eliminated FLAG-Rab40c co-precipitation with PP6. To exclude the possibility that ANKRD28 and ANKRD52 can compensate for each other, we co-depleted them using individual siRNAs. However, PP6R1 can still co-precipitate with FLAG-Rab40c in these cells (Fig. 3F). Interestingly, ANKRD28 knock-down did slightly decrease the levels of PP6R1 co-precipitating with Rab40c. Similarly, PP6R1 knock-down also diminished the amount of ANKRD28 and ANKRD52 co-precipitating with Rab40c (Fig. 3F). Taken together, this indicates that Rab40c likely interacts with both PP6R1 and ANKRD28 (and possibly ANKRD52). Collectively, we proposed that Rab40c interacts with the PP6R1/ANKRD28/PP6c complex by binding to the c-terminus of PP6R1, as well as ANKRD28 (Fig. 3G).

### Rab40 ubiquitylates and degrades ANKRD28

CRL5 complexes mediate target protein ubiquitination and subsequent degradation (Zhang & Sun, 2020, Zhao, Xiong et al., 2020). Typically, the specificity of CRL5 complex is determined by the SOCS-containing subunit that serves as an adaptor between substrate proteins and CRL5. Thus, we next examined whether Rab40c/CRL5 may regulate ANKRD28 ubiquitylation and degradation. As shown in Figure 4A-B, overall cellular levels of ANKRD28 were dramatically increased in Rab40c-KO MDA-MB-231 cells, while having little effect on the total levels of PP6c or PP6R1-3. To further confirm the increase in ANKRD28 levels is caused by depletion of Rab40c, we generated Rab40c KO cell lines stably expressing the wild-type FLAG-Rab40c or FLAG-Rab40c-4A. As shown in Figure 4C-D, reintroduction of wild-type Rab40c into the Rab40c-KO cells decreased ANKRD28 protein level, while expression of FLAG-Rab40c-4A did not have any effect on ANKRD28 (Fig. 4C-D), thus, demonstrating that ANKRD28 protein level changes are dependent on Rab40c binding to Cul5.

**Figure 4.**
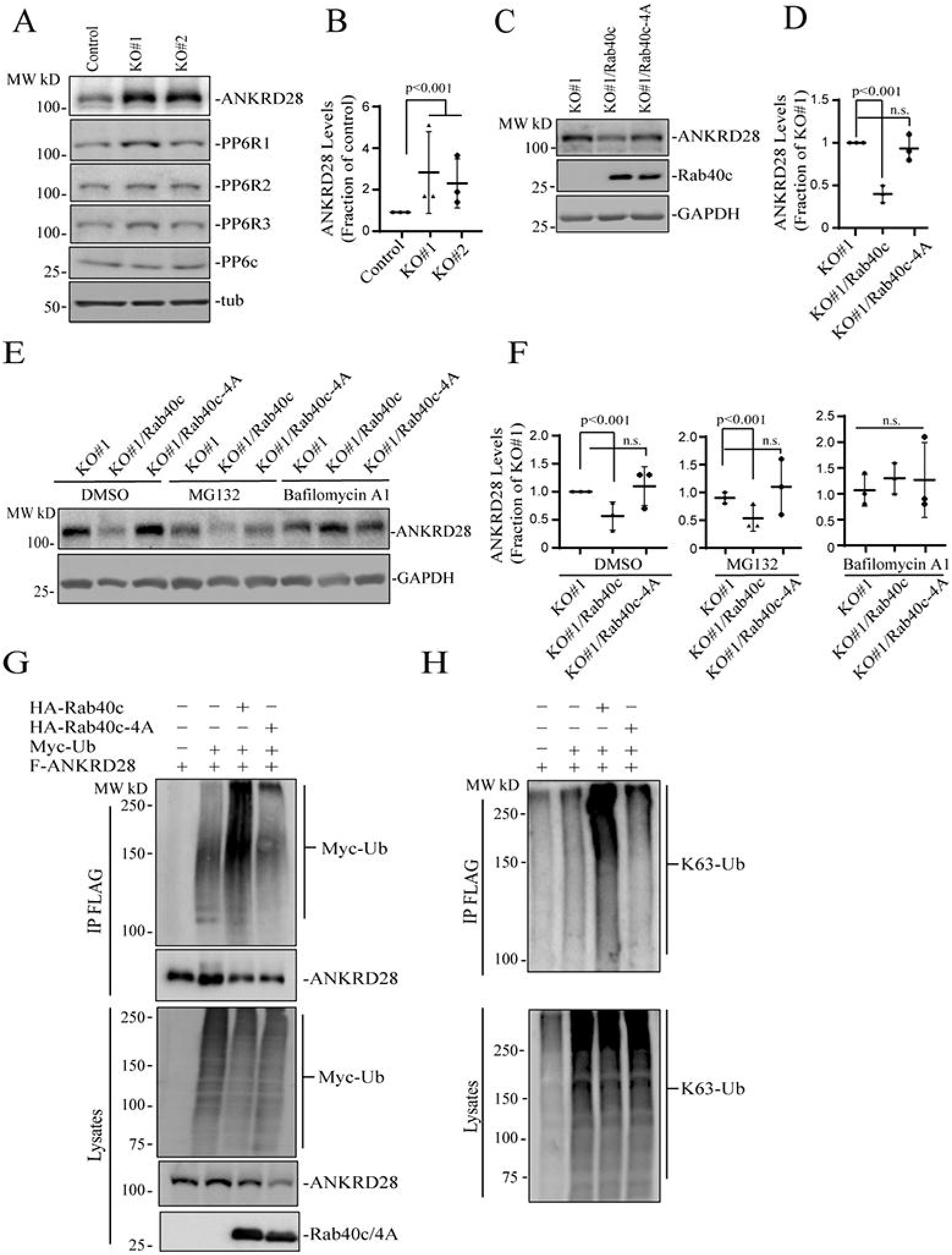
Rab40c regulates ANKRD28 ubiquitination and degradation. (A-B) Western blotting analysis of cell lysates from control and Rab40c-KO cells using indicated antibodies. Panel B shows quantification of ANKRD28 protein levels. The value shown represents the means and SEM derived from three different experiments and normalized against tubulin levels. (C-D) Western blotting analysis of cell lysates from Rab40c-KO, Rab40c-KO expressing FLAG-Rab40c or FLAG-Rab40c-4A cells using indicated antibodies. Panel D shows quantification of ANKRD28 protein levels. The value represents the means and SEM derived from three different experiments and normalized against tubulin levels. (E-F) Western blotting analysis of cell lysates from Rab40c-KO, Rab40c-KO expressing FLAG-Rab40c or FLAG-Rab40c-4A cells using anti-FLAG and anti-GAPDH antibodies. Cells were treated with DMSO, MG132 (proteosomal inhibitor), Bafilomycin A1 (lysosomal/autophagy inhibitor). (G-H) *In vivo* ANKRD28 ubiquitylation assay. 293T cells were transfected with indicated plasmids for 24 h. After treated with 100 nm Bafilomycin A1 overnight, cells were harvested and immunoprecipitated with anti-FLAG antibody followed by western blotting for either anti-Myc, anti-FLAG, anti-HA or anti-poly-Ub-K63 antibodies.

Next, we decided to investigate the cause for the increase in ANKRD28 levels. Consistent with the involvement of Rab40c-CRL5 in mediating the degradation of ANKRD28, mRNA levels of ANKRD28 in control and Rab40c KO cells, quantified by real-time qPCR, were comparable (Supplemental Fig. 3A). It is well-established that ubiquitylation targets proteins for degradation in both proteasomes (K48-Ub linkage) and lysosomes (K63-Ub linkage)(Corn & Vucic, 2014, Damgaard, 2021). Thus, we next set out to determine whether Rab40c targets ANKRD28 for proteasomal or lysosomal degradation. To that end, we treated Rab40c KO cells with either lysosomal inhibitor bafilomycin A1 or proteasomal inhibitor MG132. Surprisingly, bafilomycin A1, but not the MG132 treatment, rescued Rab40c-induced increase in ANKRD28 protein level (Fig.4E), suggesting that Rab40c/CRL5 mediate lysosomal degradation of ANKRD28.

Next, we asked whether ANKRD28 can be directly ubiquitylated by Rab40c/CRL5. To examine this, we first transfected 293T cells with FLAG-ANKRD28, Myc-Ub, HA-Rab40c, and HA-Rab40c-4A individually or in combinations (Fig.4G). Lysates were then immunoprecipitated with anti-FLAG antibodies and blotted for Myc-Ub with anti-Myc antibodies. When Myc-Ub was co-transfected with FLAG-ANKRD28 in the presence of the lysosomal inhibitor bafilomycin A1, the high molecular weight species were detected (presumably ubiquitylated ANKRD28), which were significantly enhanced by co-transfecting HA-Rab40c. Importantly, co-transfection of HA-Rab40c-4A with FLAG-ANKRD28 did not increase ANKRD28 polyubiquitylation, supporting that Rab40c mediates ANKRD28 ubiquitylation in a Cul5-depedent manner (Fig. 4G).

Our data so far suggests that Rab40c/CRL5-dependent polyubiquitylation may target ANKRD28 to lysosomes for degradation, which is usually mediated by K63-linked polyubiquitylation. Consistent with this, Western blot analysis with anti-K63-specific anti-ubiquitin antibodies showed that Rab40c/CRL5 increases K63-linked polyubiquitylation of ANKRD28 (Fig. 4G). Taken together, these results support our hypothesis that Rab40c forms a Cul5-based-ubiquitin E3 ligase complex to ubiquitylate ANKRD28 and promote its lysosomal degradation.

### ANKRD28 regulates FA formation

ANKRD28 is a large scaffolding protein with 26 ankyrin repeats which has been reported to regulate cell migration (Kiyokawa & Matsuda, 2009, Tachibana, Kiyokawa et al., 2009). We therefore examined whether ANKRD28 is present at the lamellipodia where it may also regulate FAs. To that end, we first generated MDA-MB-231 cell lines stably expressing FLAG-tagged ANKRD28, or its binding partner PP6R1 (Supplemental Fig. 3B), to examine their localization by immunofluorescence microscopy. As shown in Figure 5A, the majority of ANKRD28 protein was localized in the cytosol, but a fraction of FLAG-ANKRD28 could be observed at the lamellipodia, where it colocalizes with actin ruffles. Similarly, a sub-population of FLAG-PP6R1 could also be observed at the leading edge of the lamellipodia (Fig. 5B), suggesting that PP6 complexes containing ANKRD28 and PP6R1 may function at the leading edge of the migrating cell, although additional experiments will be needed to further demonstrate that.

**Figure 5.**
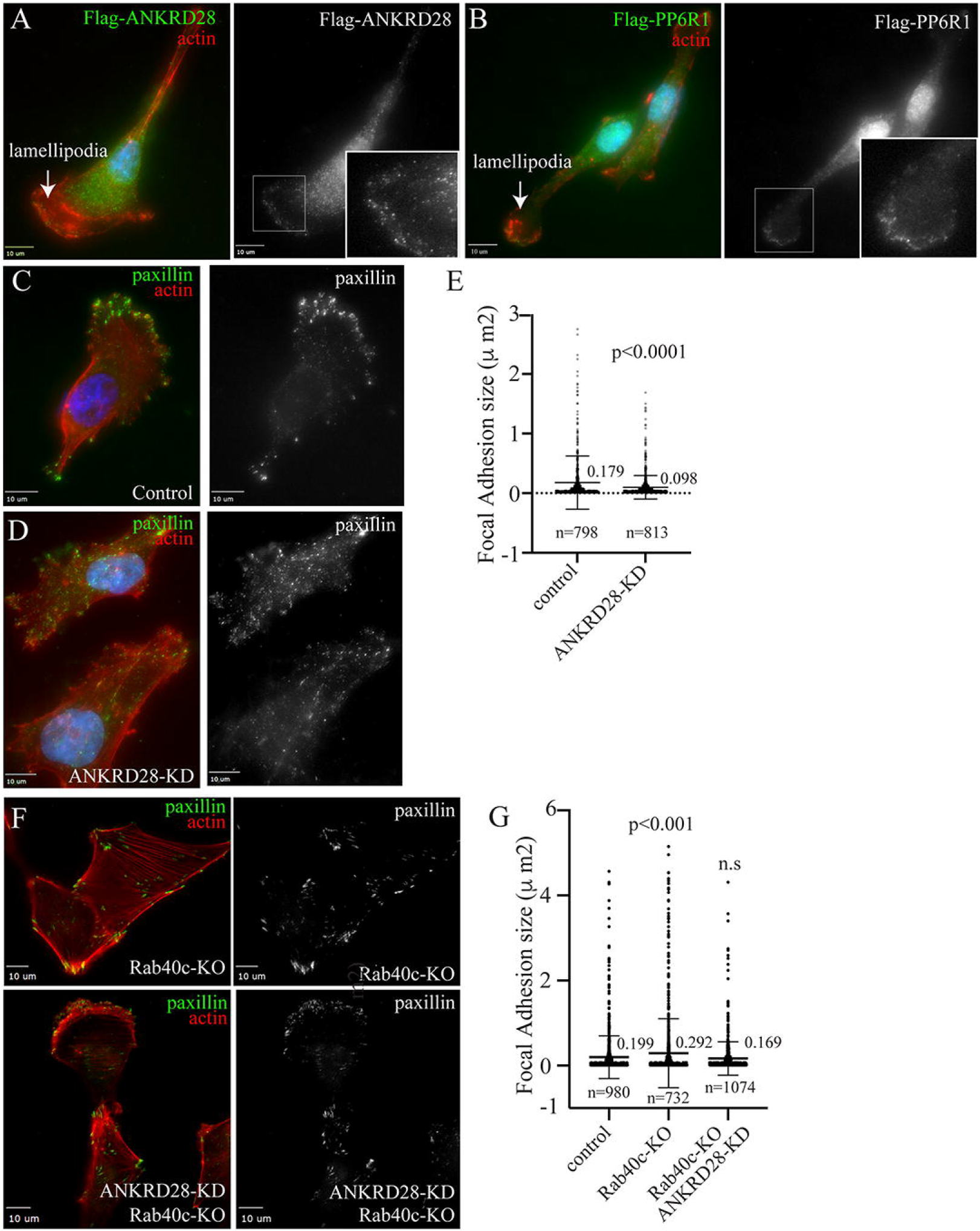
ANKRD28 regulates FA formation. (A-B) MDA-MB-231 cells stably expressing FLAG-ANKRD28 or FLAG-PP6R1 were plated on collagen-coated coverslips and then fixed and stained with anti-FLAG antibodies (green) and phalloidin-AlexaFluor594 (red). Inset regions of interest highlight ANKRD28 or PP6R1 positive puncta on the leading edge. Arrows point to the leading edge of lamellipodia. (C-E) MDA-MB-231 cells were transfected with control or ANKRD28 siRNA. Cells were then plated on collagen-coated coverslips, fixed and stained with anti-paxillin antibody (green) and phalloidin-AlexaFluor594 (red). Panel E shows quantification of FA size. n = number of FAs analyzed. Shown data are the means and standard deviations derived from three independent experiments. (F-G) control or Rab40c-KO cells were transfected with ANKRD28 siRNA and then plated on collagen-coated coverslips and stained with anti-paxillin antibody (green) and phalloidin-AlexaFluor594 (red). Panel G shows quantification of FA size in control, Rab40c-KO, and Rab40c-KO plus ANKRD28 siRNA. n = number of FAs analyzed. Shown data are the means and standard deviations derived from three independent experiments.

To test whether ANKRD28 regulates FA formation at the leading edge of the cell, we next used siRNA to knock-down ANKRD28 in MDA-MB-231 cells (Supplemental Fig. 3C), and then cells were stained with an anti-paxillin antibody to visualize the size and distribution of FAs. As expected, in control cells FAs were mostly present at the leading-edge of lamellipodia (Figure 5C). In contrast, in ANKRD28-depleted cells, FAs were smaller in size and situated not only at the periphery of the leading edge but scattered throughout the entire cell (Fig. 5C-F). Importantly, the ANKRD28 knock-down phenotype is opposite to what was observed in Rab40c-KO cells, which generate bigger FAs. That is consistent with our hypothesis that the Rab40c/CRL5 complex regulates ANKRD28 degradation and inactivation of an ANKRD28-containing PP6 complex.

If Rab40c regulates FAs via an ANKRD28-containing PP6 complex, then ANKRD28 knockdown would be expected to reverse Rab40c-KO induced increase in FA size. To determine that, we used siRNA to knockdown ANKRD28 in Rab40c-KO cells. These cells were then fixed and stained with anti-paxillin antibodies to analyze FAs. As shown in Figure 5G-H, ANKRD28 knockdown did decreased FA size in Rab40c-KO cells. This is consistent with the hypothesis that Rab40c affects FA formation by regulating ANKRD28 degradation and activity of ANKRD28-containing PP6 complexes at the lamellipodia of migrating cells.

### Rab40c and ANKRD28 regulate FAK and Hippo signaling pathways during cell migration

Our data so far suggests that Rab40c regulates FA formation during cell migration. This regulation is presumably mediated by ANKRD28/PP6-dependent de-phosphorylation of specific target proteins that regulate FAs. Thus, next we set out to identify the identity of the target proteins that are directly or indirectly regulated by the ANKRD28/PP6 complex. One of the well-established key regulators of FA dynamics is focal adhesion kinase (FAK) (Katoh, 2020, Tapial Martinez et al., 2020). Importantly, FAK is tightly regulated by several tyrosine and serine/threonine kinases, thus, could be a candidate for ANKRD28/PP6-dependent de-phosphorylation. Among several Ser/Thr phosphorylation sites, Ser910 has emerged as one of the key regulators of FAs (Villa-Moruzzi, 2007, Zheng, Xia et al., 2009). Specifically, phosphorylation of FAK-Ser910 was shown to increase FA turnover, presumably by regulating recruitment of paxillin and several tyrosine phosphatases that then de-phosphorylate Y392. To test whether Rab40c may regulate FAs by inhibiting ANKRD28/PP6-dependent Ser910 de-phosphorylation, we have compared the levels of pFAK-Ser910 in control and Rab40c-KO cells. As shown in Supplemental Figure 4, Rab40c depletion did decrease the levels of pFAK-Ser910. That, at least in part, would contribute to the increase in FA size and number in Rab40c-KO cells (Fig. 1), although further studies will be needed to determine whether ANKRD28/PP6 directly de-phosphorylates pFAK-Ser910.

Recently, the Hippo-signaling pathway was reported to play an important role in mechanosensing ECM stiffness and regulating FAs dynamics (Mason, Collins et al., 2019, Nardone et al., 2017, Wang et al., 2021). Importantly, ANKRD28 has been suggested to bind MOB1, the key regulators of the Hippo pathway (Fig. 6A) (Couzens, Knight et al., 2013, Ohama, 2019, Rusin, Schlosser et al., 2015). However, whether ANKRD28 actually regulates Hippo-signaling has not been explored, and our data raises a very interesting possibility that Rab40c regulates Hippo signaling at the FAs through inactivation of ANKRD28/PP6 complex. MOB1 functions as an activator of Lats1 (Figure 6A), and MOB1 phosphorylation by MST1/2 increases its ability to activate Lats1 (Delgado, Carmona et al., 2020, Duhart & Raftery, 2020, Gundogdu & Hergovich, 2019, Ni, Zheng et al., 2015). Consistent with the possibility that Rab40c may regulate the Hippo pathway, we found the phosphorylation of MOB1 at Thr35 was significantly decreased, whereas the total protein level significantly increased in Rab40c-KO cells (Fig. 6B and C). Furthermore, reintroduction of wild-type Rab40c, but not Rab40c-4A mutant, into the Rab40c-KO cells rescued MOB1 phosphorylation defects and decreased the total protein level of MOB1, suggesting the changes in MOB1 phosphorylation and total protein levels may be mediated by Rab40c/CRL5 complexes (Figure 6D and E). To further confirm that changes in MOB1 phosphorylation in Rab40c KO cells are mediated by ANKRD28, we used two different siRNAs to knock-down ANKRD28. As shown in Figure 6F and G in control cells, knock-down of ANKRD28 increased the levels of pMOB1-T35, again supporting the hypothesis that the ANKRD28/PP6 complex regulates MOB1-T35 phosphorylation. Collectively, this data demonstrates that Rab40c may regulate MOB1 phosphorylation through regulating ANKRD28 ubiquitylation and degradation, thus inhibiting ANKRD28/PP6 complex activity.

**Figure 6.**
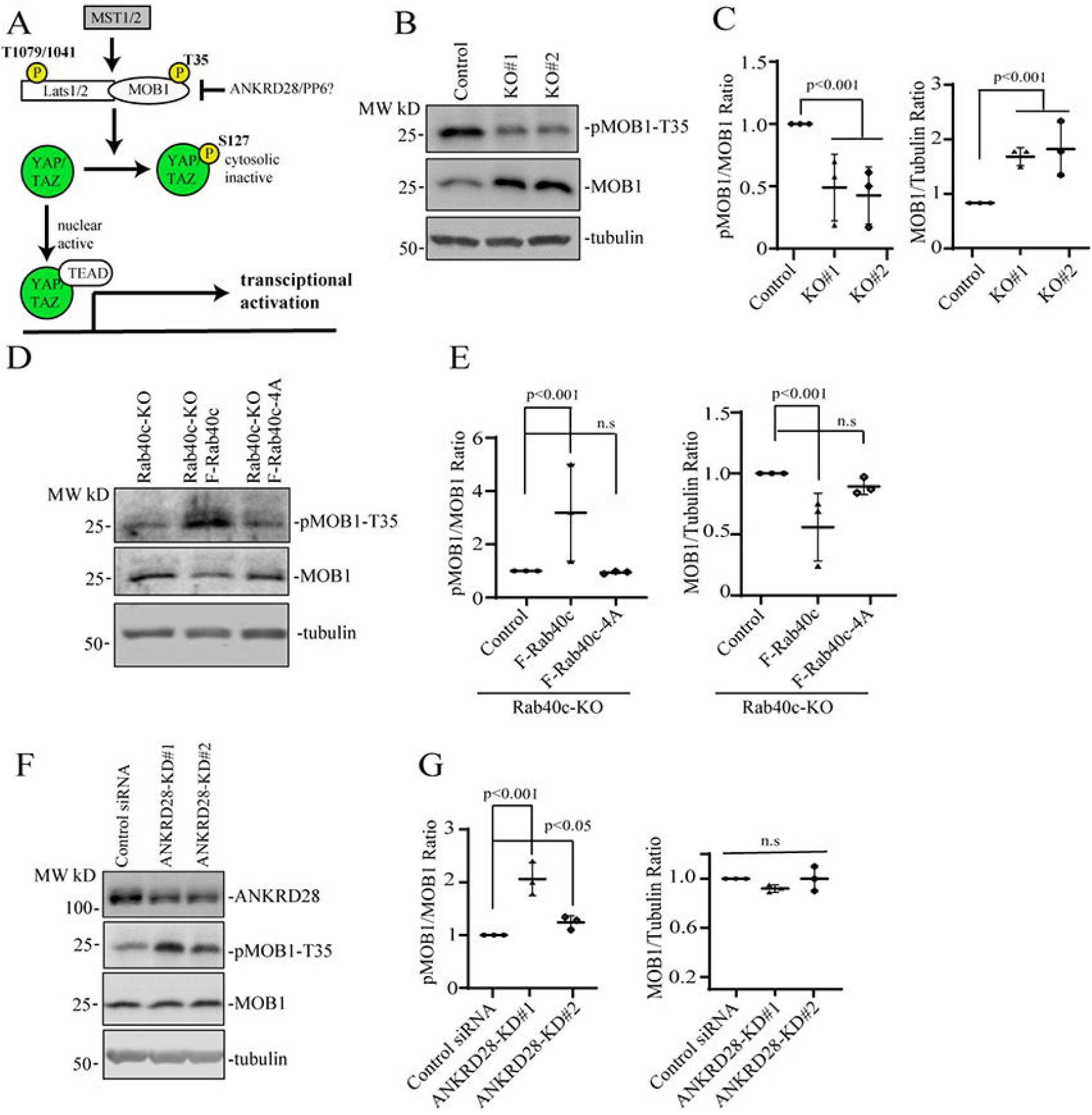
Rab40c and ANKRD28 regulates MOB1 phosphorylation. (A) A model showing Hippo-signaling pathway and a potential target of ANKRD28/PP6 complex. (B-C) Immunoblotting of cell lysates from control and Rab40c-KO cells with anti-pMOB1-T89, anti-MOB1 and anti-tubulin antibodies. Panel C shows quantification of pMOB1/MOB1 and MOB1/tubulin ratio. The data shown represents the means and SEM derived from three different experiments and normalized against tubulin levels. (D-E) Immunoblotting of cell lysates from Rab40c-KO, Rab40c-KO expressing FLAG-Rb40c or FLAG-Rab40c-4A cells with anti-pMOB1-T35, anti-MOB1 and anti-tubulin antibodies. Panel E shows quantification of pMOB1/MOB1 and MOB1/tubulin ratio in (D). The data shown represents the means and SEM derived from three different experiments and normalized against tubulin levels. (F-G) MDA-MB-231 cells were transfected with ANKRD28 siRNA or non-targeting control. Cell lysates were then immunoblotted with indicated antibodies. Panel G shows quantification of pMOB1/MOB1 and MOB1/tubulin ratio. The shown data represents the means and SEM derived from three different experiments and normalized against tubulin levels.

Our findings that Rab40c regulates MOB1 protein levels and phosphorylation prompted us to test whether Hippo downstream transcription factors YAP and TAZ are also activated, because LATS1/MOB1 complex mediates YAP/TAZ phosphorylation (Battilana, Zanconato et al., 2021, Meng, Moroishi et al., 2016). Phosphorylation of YAP/TAZ acts as a negative regulator of YAP/TAZ by keeping them in cytosol and away from the nucleus (Kwon, Kim et al., 2021, Rausch & Hansen, 2020). Due to a significant similarity between YAP and TAZ in their sequences, it is challenging to find highly specific YAP antibodies, thus, we focused on TAZ. Given that nuclear-cytoplasmic localization reflects TAZ activity, we used immunofluorescence staining to examine the localization of TAZ. As shown in Figure 7A-C, Rab40c-KO increased TAZ translocation from the cytoplasm into the nucleus (Figure 7C). The increase seems to be Rab40c-KO specific because neither Rab40a nor Rab40b knock-out led to a similar increase in nuclear TAZ (Fig. 7D) and knocking out all three Rab40 isoforms (TKO) did not further increase the nuclear TAZ as compared to Rab40c-KO cells (Figure 7D).

**Figure 7.**
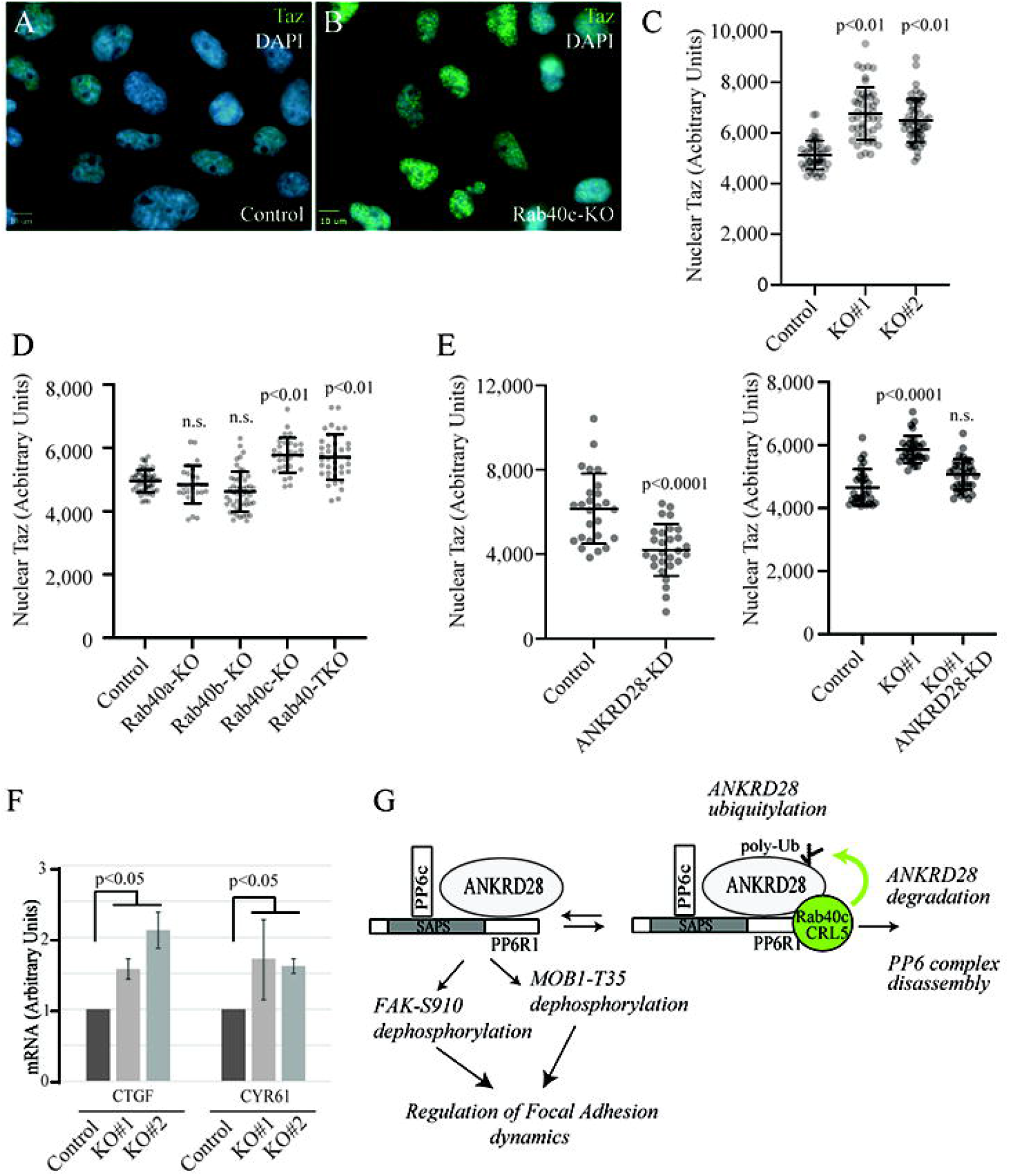
Rab40c regulates Hippo signaling in MDA-MB-231 cells. (A-C) Control and Rab40c-KO MDA-MB-231 cells were plated on collagen-coated coverslips. Cells were then fixed and stained with anti-TAZ antibody (green) and DAPI (blue). Panel C shows quantification of nuclear TAZ fluorescence intensity. Data shown are the means and standard deviations derived from three independent experiments. (D) Quantification of nuclear TAZ fluorescence intensity in control, Rab40c-KO, Rab40b-KO, Rab40c-KO, and Rab40-TKO cells stained as described in (A). (E) Quantification of nuclear TAZ fluorescence intensity in MDA-MB-231 cells transfected with ANKRD28 siRNA or non-targeting control (left panels). Quantification of nuclear TAZ fluorescence intensity in control or Rab40c-KO cells transfected with either non-targeting control siRNA or Rab40c ANKRD28 siRNA (right panels). (F) qPCR analysis of CTGF and CYR61 mRNAs (known Hippo pathway targets) in control and Rab40c-KO MDA-MB-231 cells. (G) Proposed model for Rab40c function in regulation FAs dynamics. The PP6 complex containing ANKRD28 and PP6R1 dephosphorylates FAK-S910 and/or pMOB1-T35 to regulate FAs dynamics. When Rab40c binds to PP6 through ANKRD28 and PP6R1, it leads to ANKRD28 ubiquitylation and subsequent degradation which results in PP6 disassembly and inactivation.

Since Rab40c/CRL5 mediates ubiquitylation-dependent degradation of ANKRD28, we next examined whether ANKRD28 is a positive TAZ regulator, and whether ANKRD28 knockdown inhibits TAZ nuclear localization. As shown in Figure 7E, knockdown of ANKRD28 by siRNA significantly reduced nuclear TAZ in MDA-MB-231 cells. Importantly, knockdown of ANKRD28 in Rab40c KO cells reversed Rab40c-KO induced activation of TAZ (Figure 7E). Taken together, our data is consistent with the hypothesis that Rab40c-KO promotes TAZ nuclear localization by increasing ANKRD28/PP6-dependent de-phosphorylation of Hippo signaling pathways regulators, such as MOB1.

Nuclear TAZ interacts with transcription factor TEAD to operate as a coactivator to increase transcription of their target genes such as CTGF and CYR61. If Rab40c KO leads to an increase in nuclear TAZ, one would predict that Rab40c KO should also increase the transcription of TAZ target genes. Consistent with this, we found the mRNA expression levels of CTGF and CYR61 were higher in Rab40c-KO than in the control cells, as determined by qPCR analysis (Fig. 7F). Altogether, our data supports that Rab40c forms a Cul5-based ubiquitin E3 ligase to ubiquitylate and degrade ANKRD28. This then results in inhibiting ANKRD28/PP6 complex activity and subsequent increases in phosphorylation of several important signaling molecules including FAK and MOB1, thus, regulating FAs dynamics during cell migration.

## DISCUSSION

Rab40c belongs to a unique Rab40 subfamily of small monomeric GTPases that contain SOCS domain at their C-terminus, thus, binding Cul5 to form a ubiquitin E3 ligase complex (Rab40/CRL5) to control protein ubiquitylation and degradation (Duncan et al., 2021a). All vertebrates express two Rab40 family isoforms, Rab40b and Rab40c. While Rab40b and Rab40c are closely related isoforms, it has been suggested that they may ubiquitylate and regulate different, but probably overlapping, subsets of target proteins. Indeed, RACK1 and Varp, the only two proposed Rab40c/CRL5 target proteins, do not appear to be ubiquitylated by Rab40b (Day, Whiteley et al., 2018, Linklater et al., 2021, Yatsu, Shimada et al., 2015), while Rap2 GTPase appears to be ubiquitylated predominately by Rab40b (Duncan et al., 2021b). Thus, determining the repertory of Rab40b and Rab40c-specific target proteins will be the key to understanding functions of the Rab40 subfamily of small monomeric GTPases, and is one of the goals of this study.

Regulating cell migration recently emerged as one of the major functions of the Rab40 subfamily of GTPases. It has been shown that Rab40b/CRL5 regulates cell motility and invasion by ubiquitylation of EPLIN (Linklater et al., 2021), as well as by mediating MMP2 and MMP9 transport to the invadopodia (Jacob et al., 2013). Similarly, Rab40a, an isoform expressed only in higher primates, regulates degradation of RhoU (Dart et al., 2015), one of the regulators of cell adhesion to extracellular matrix (ECM). However, it remains unclear whether Rab40c is also involved in regulating cell motility, and what proteins could be its targets of ubiquitylation. In this study, we show that Rab40c plays an important role in controlling focal adhesion (FA) formation during cell migration, since Rab40c depletion significantly increases the number and size of FAs, while also disrupting polarization of FA formation at the leading edge of lamellipodia. Furthermore, while Rab40c previously was suggested to regulate lipid droplet formation (Luo, Li et al., 2017, Tan, Wang et al., 2013), we did not find any evidence that Rab40c plays a similar role in migrating MDA-MB-231 cells. Thus, taken together, all this data suggests that all Rab40 subfamily isoforms regulate, at least in part, actin and FA dynamics during cell migration by ubiquitylating and regulating isoform-specific target proteins.

Rab proteins are key regulators of intracellular membrane trafficking, and each Rab protein has a distinct location corresponding to its functions. Here we show that Rab40c localizes at two specific compartments: Golgi, and the leading edge of the lamellipodia, implying that Rab40c may regulate membrane trafficking from the Golgi to the cell surface, as well as plasma membrane and actin dynamics during migration. In line with this, we found that Rab40c-KO cells form more, and bigger FAs, which evenly distribute throughout the cell. FAs are integrin-containing, multi-protein structures that link the intracellular cytoskeleton to the ECM. The number and localization of FAs are tightly controlled during cell migration, with coordinated assembly and turnover of FAs at the front of the migrating cell body, and disassembly at the rear. Intriguingly, Rab18, an endoplasmic reticulum (ER)-resident protein, regulates kinectin-1 transport toward the cell surface to form ER– FA contacts, thus promoting FA growth during cell migration. Rab18 is closely related to the Rab40 family, and it has been suggested that Rab40 family proteins are expanded from ancestral Rab18 during metazoan evolution. Thus, it is not surprising that they share a partially redundant role in some cellular processes.

The Rab40 subfamily of GTPases have a conserved SOCS domain at their C-terminus, and it has been shown that Rab40b binds directly to Cul5 and its accessory proteins ElonginB/C (Duncan et al., 2021b, Linklater et al., 2021). Here we show that human Rab40c also binds Cul5 and ElonginB/C, and that this binding is blocked by mutating a Cul-box within the SOCS domain. We also identified several putative Rab40c-interacting proteins, including protein phosphatase 6 (PP6) complex. PP6 is heterotrimeric complex that belongs to the serine/threonine phosphatase family, comprising of a single catalytic subunit (PP6c), one of a PP6 regulatory subunit (PP6R1,2 or 3), and one of an ankyrin repeat domain scaffolding subunit (ANKRD28, 44 or 52) (Ohama, 2019, Stefansson et al., 2008). PP6 complex plays an important role in many fundamental cellular processes, but the regulatory mechanism for PP6 complex activity remains largely unknown. Since the catalytic PP6 subunit appears to be quite promiscuous, it has been proposed that the specificity of PP6 activity is regulated by its scaffolding (ANKRD28, 44, 52) and regulatory (PP6R1-3) subunits. Thus, it has been hypothesized that specific ANKRD and PP6R subunits define spatiotemporal properties of PP6 complex activity and determine the target protein specificity. However, so far, we know very little about specific functions of different ANKRD and PP6R subunits. In this study, we found that Rab40c interacts with the PP6 complex, which leads to ubiquitylation and degradation of its scaffolding subunit ANKRD28. Though which subunit is responsible for directly recruiting Rab40c to the PP6 complex needs to be further assessed, our data suggests that Rab40c co-binds to both ANKRD28 and PP6R1, likely recruiting Rab40c to the assembled PP6 complex. Importantly, regulation of ANKRD28 by Rab40c is Cul5 dependent because Rab40c, but not Rab40c-4A mutants, can enhance ANKRD28 ubiquitylation.

Since, in addition to plasma membrane, Rab40c also localizes to the Golgi, we examined the subcellular localization of ANKRD28 and PP6R1. However, we did not observe ANKRD28 or PP6R1 at the Golgi, but instead we found that both these proteins are present at the leading edge of lamellipodia. Consequently, we hypothesize that Rab40c/CRL5 may regulate FAs through induction of localized ANKRD28 degradation that would lead to localized disassembly and inactivation of ANKRD28 and PP6R1-containing PP6 complexes at the leading edge of lamellipodia. In fact, ANKRD28 has been previously implicated in regulation of FA dynamics and cell migration (Kiyokawa & Matsuda, 2009, Tachibana et al., 2009), although how ANKRD28 affects FA dynamics remains largely unclear. In this study we found that ANKRD28 knockdown leads to a decrease in FA size and a loss of polarized FA distribution at the leading edge of the migrating cell. Importantly, depletion of ANKRD28 in Rab40c KO cells partially restored both the size and distribution of FAs, suggesting that the Rab40c-KO phenotype was, at least partially, due to an increase in ANKRD28 protein level and presumably over-activation of PP6 complex at the leading edge.

ANKRD28 is an essential scaffolding component of the PP6 complex, thus, we predicted that Rab40c-dependent ubiquitylation and degradation of ANKRD28 leads to localized inactivation of a specific subset of PP6 (PP6R1/ANKRD28/PP6c), in turn controlling the phosphorylation of some signaling molecules important for FAs dynamics. It is well established that FAK-Src signaling plays a crucial role in regulating FAs dynamics. Phosphorylation of FAK-S910, which is mediated by ERK, promotes FAK dephosphorylation at Y397, and is necessary for invasive cell migration (Zheng et al., 2009). We found pFAK-S910 levels are decreased in Rab40c-KO cells, while total FAK remained constant. That raises an intriguing possibility that ANKRD28/PP6 directly dephosphorylates pFAK-S910. Another possibility is that ERK activity is decreased in Rab40c-KO cells because PP6 has recently been identified as a key negative regulator of ERK signaling by dephosphorylating MEK (Cho, Lou et al., 2021). Thus, while additional studies will be needed to determine the mechanisms of PP6-dependent FAK regulation, our data implies that the Rab40c-ANKRD28/PP6 pathway contributes to regulating FAK activity and FA disassembly during cancer cell migration (Fig. 7G).

PP6 complex was also identified in interactome analysis of the Hippo signaling pathway, and it was proposed that PP6 may be a regulator of Hippo-signaling (Couzens et al., 2013, Ohama, 2019, Sarmasti Emami, Zhang et al., 2020). However, the precise role of PP6 in the Hippo pathway remains unknown. Importantly, interactions between the Hippo signaling pathway components YAP/TAZ and FAs have been revealed recently (Mason et al., 2019, Nardone et al., 2017, Wang et al., 2021). It was shown that FAs act as a hub for sensing and transmission of mechanical cues to regulate YAP/TAZ activation. In turn, YAP/TAZ regulate cell mechanics by controlling FA assembly through co-transcription of genes encoding for various FA regulators. Based on this data, we examined the possibility that Rab40c-ANKRD28/PP6 directly regulates Hippo-signaling to control FAs dynamics. Consistent with this hypothesis, pMOB1-T35 levels decreased in Rab40c KO cells. In contrast, knocking down ANKRD28 increased pMOB1-T35. Finally, Rab40c KO induced decrease in MOB1 phosphorylation could be rescued by an overexpression of wild-type, but not Cul5-binding mutants, of Rab40c.

MOB1 is a key regulator of large tumor suppressor 1/2 (Lats1/2) kinases, and phosphorylation of pMOB1-T35 promotes its binding to Lats1/2 (Couzens et al., 2013, Meng et al., 2016). Phosphorylation of YAP and TAZ by Lats/MOB1 kinase complex results in YAP/TAZ cytoplasmic retention and inhibits their transcriptional activities. Consistent with this, we confirmed that Hippo-signaling is affected in Rab40c-KO cells by TAZ immunostaining and qPCR quantification of YAP/TAZ target genes. More interestingly, a recent study showed that Rab40c is downregulated upon YAP stimulation (Moon, Lee et al., 2020), thus, the Rab40c-PP6 and YAP/TAZ may constitute a feed-forward loop to regulate FAs dynamics in migrating cells. Taken together, our data suggests that the Cul5/Rab40c-ANKRD28/PP6 axis is an important regulator of Hippo-signaling and FAs dynamics (Fig. 7G).

Although our data demonstrate that Rab40c-ANKRD28/PP6 affects FAs formation by co-regulating FAK and Hippo-YAP/TAZ signaling (Figure 7G), many questions remain to be addressed in the future. For example, it remains completely unknown what regulates formation and activity of the Rab40c/CRL5 complex, and whether this complex has distinct functions at the plasma membrane and Golgi. Can Rab40c-PP6 regulate vesicles (like MMP2/9-containing secretory vesicle) trafficking? Additionally, it is becoming clear that Rab40/CRL5 complexes regulate ubiquitylation of multiple proteins, thus, what are other Rab40c/CRL5 targets and what are their functions? Finally, do our findings have any clinical implications? Given about 10% of melanoma patients harbor PP6c inactivating mutations, and dysregulated Hippo pathways are also associated with various diseases, especially with cancer, answering these questions will be very interesting and be the focus of future studies.

## MATERIALS AND METHODS

### Cell Culture and Transfection

All cell lines were cultured as described previously ((Jacob et al., 2013). Briefly, human embryonic kidney (HEK) 293T cells were grown in complete Dulbecco’s modified Eagle medium (DMEM supplemented with 10% fetal bovine serum and 100 μg/ml of penicillin and streptomycin) at 37 °C in a 5% CO2 atmosphere. MDA-MB-231 cells were grown in complete DMEM supplemented with 1μg/ml human recombinant insulin, 1% non-essential amino acids, and 1% sodium pyruvate. Cell lines were routinely tested for mycoplasma. All cell lines used in this study were authenticated and are in accordance with American Type Culture Collection standards. 293T cells were grown to 70–80% confluence and transfected using the Lipofectamine 2000 (Invitrogen) transfection reagent according to the manufacturer’s instructions. MDA-MB-231 cells were grown to 80-90% confluence and transfected using JetPRIME (polyplus). Lipofectamine RNAiMAX (Invitrogen) was used for transfection of siRNAs both in 293T and MDA-MB-231 cells.

### Mammalian Expression Constructs

Human Rab40a, Rab4-b, and rRb40c plasmids were purchased from the Functional Genomics Core Facility at the University of Colorado. Human ANKRD28/44/52, PP6R1/2/3, PP6c, and Myc-Ub were described previously (Couzens et al., 2013, Han, Foster et al., 2012). Expression plasmids of GFP-Rab40c, FLAG-Rab40c, HA-Rab40c, and FLAG-PP6R1 deletion mutants were constructed by PCR, followed by subcloning into the pRK7 or pGPS vector containing an N-terminal FLAG or HA tag. FLAG-Rab40c-4A mutant was generated by PCR using the following primers (Integrated DNA Technologies): Forward: GTCGTCGACATGGGCTCGCAGGGCAGTCCGGTG, Reverse: TGCAGCCTTGTCGATGAGG and Forward: GCGGCCGTCACCATCAAG, Reverse: TAGCGGCCGCTAGGAGATCTTGCAGTTAC. All plasmids were validated by DNA sequencing.

### Antibodies

The following antibodies were used in this study: anti-FLAG (clone M2, WB 1:1,000, Sigma), anti-GAPDH (WB 1:5,000, UBPBio), FAK S910 (WB 1:1,000, 44-596G, Invitrogen), total FAK (WB 1:1,000, 610087, BD Biosciences), paxillin (IF 1:500 Transduction labs). Anti-HA (WB 1:500, IP 2 µg/1 mg cell lysate, SC F-7), anti–α-tubulin (WB 1:5,000, 23948), anti-Rab40c (WB 1:500, H-8 sc514826), cul-5 (WB 1:500, H-300), and mouse ANKRD28 (WB 1:500) were purchased from Santa Cruz Biotechnology. MOB1(E1N9D) and p-MOB1(D2F10) were purchased from cell signaling technology. Rabbit anti-SAPS1/2/3, ANKRD28/52, PP6c were purchased from Bethyl Laboratories.

### Identification of Rab40c-interacting Protein

A lentivirus-based method was used to generate stable cell lines expressing FLAG-Rab40c or FLAG-Rab40c-4A as described previously ((Han et al., 2012). Putative Rab40c-binding proteins were identified by coimmunoprecipitation using anti-FLAG antibody–coated beads as described previously (Prekeris, Klumperman et al., 2000). Briefly, 50 µg affinity purified anti-FLAG antibody was bound to 100 µl Protein G–Sepharose beads. Antibodies were then cross-linked to beads using dimethyl pimelimidate dihydrochloride. Anti-FLAG antibody beads were then incubated with 2 ml of 1 mg/ml Triton X100 cellular lysates (PBS, 1% Triton X-100, and 10 mM PMSF), followed by a wash with 5 ml PBS. Proteins were eluted from anti-FLAG antibody beads with 1% SDS and then analyzed using tandem mass spectrometry (Proteomics Core on campus). The UniProtKB/SwissProt human database was used for protein identification. Nonspecific contaminants were identified and eliminated by the following criteria: (1) presence in IgG control; (2) presence in the CRAPome database and (3) all RNA, DNA binding proteins, and mitochondrial proteins were considered a contaminant.

### CRISPR-Cas9 knockout lines

Guide RNAs for Rab40c targeting 59-TACCGTTACTGTAGGCGTAC-39 (exon 1) and 59-AGGTAGTCGTAGCTCTTCAC-39 (exon 3) were transfected into MDA-MB231 cell line with Dox-induced Cas9. Cells were split 24 h after transfection and seeded for single colonies and then were screened by Western Blotting, followed by PCR cloning and genotyping. Rab40a, b or triple knockout lines has been described previously (Linklater et al., 2021). For each knockout line, two different clones were used for all experiments.

### Quantitative PCR (qPCR)

Total RNA was extracted using TRIzol (Invitrogen) according to the manufacturer’s protocol. Reverse transcription to cDNA was performed with SuperScript III (Invitrogen) using random hexamer primers. qPCR was performed using iTaq SYBR Green qPCR Master Mix on Applied Biosystems ViiA7 Real Time PCR System. The qPCR amplification conditions were 50°C (2 min), 95°C (10 min), 40 cycles at 95°C (15 s), and 60°C (1 min). Targets were normalized to GAPDH. The following primers used for qPCR: ANKRD28 forward: ACTGCTCTCCACGGTAGATTC and reverse: GGGGAACATTCCATGTATGCC; CTGF forward: ACCGACTGGAAGACACGTTTG and reverse: CCAGGTCAGCTTCGCAAGG; CYR61 forward: AGCCTCGCATCCTATACAACC and reverse: TTCTTTCACAAGGCGGCACTC; GAPDH forward: CTGGGCTACACTGAGCACC and reverse: AAGTGGTCGTTGAGGGCAATG.

### Immunoprecipitation and Immunoblotting Assays

For non-denaturing immunoprecipitation, cells in a 100-mm dish were harvested and washed with 1× PBS, then lysed with 1.0 ml ice-cold cell lysis buffer (20 mM Tris-HCl, pH 7.6, 150 mM NaCl, 2 mM EDTA, 1% Triton X-100, 10% glycerol) with protease inhibitor cocktails (Roche). After clearing lysates by centrifugation, supernatants were incubated with 2μg of an appropriate antibody or control IgG for 4 hr at 4°C, then supplemented with 50 μl protein G beads. After overnight rocking, protein G beads were pelleted by centrifugation and washed three times with the cell lysis buffer plus 0.5 M NaCl. Bound proteins were eluted in 50 μl 1× SDS sample buffer.

For denaturing immunoprecipitation (used in ubiquitylation assays), cells in a 100-mm dish were lysed in 1 ml cell lysis buffer plus 1% SDS. Cell lysates were collected and then heated at 95°C for 15 min. After centrifugation, supernatants were diluted with the cell lysis buffer to reduce SDS concentration to 0.2%. The immunoprecipitation assay was performed as described above, except that 5 μg anti-FLAG antibody was used in each reaction. Eluates (40 μl) were resolved in SDS-PAGE and transferred to nitrocellulose membranes for immunoblotting assays. Immunoblotting images were captured using a ChemiDoc MP Imaging system (Bio-Rad).

### Ubiquitylation Assay

Ubiquitylation assay was performed as described previously (Linklater et al., 2021). Briefly, 293T cells (∼80% confluency) were transfected with plasmids expressing pRK5-FLAG-ANKRD28 with or without pRK5-Myc-Ub, pRK7-HA-Rab40c or pRK7-HA-Rab40c-4A using Lipofectamine 2000. After 24hrs, cells were treated with 100nM Bafilomycin-A1 (Selleckchem S1413) overnight. Then, cells were lysed in 1% SDS for denaturing immunoprecipitation as described above. Bound proteins were eluted in 50µl 1X SDS sample buffer. Eluates (20µl) were resolved via SDS-PAGE and transferred to nitrocellulose membranes for immunoblotting. Blot images were captured using a ChemiDoc MP Imaging system (Bio-Rad).

### siRNA knockdown

For ANKRD28/52 and PP6R1 knockdown, siRNAs were purchased from Sigma. Mission siRNA universal negative control (SIC001; Sigma), ANKRD28 siRNAs (SASI_Hs01-00173856) and (SASI_Hs01_00173857), ANKRD52(SASI_Hs02_00368435) and PP6R1(SASI_Hs01_00222781) were transfected using Lipofectamine RNAiMAX (Invitrogen) according to manufacturer protocol.

### Immunofluorescent microscopy

MDA-MB-231 cells were seeded onto collagen-coated glass coverslips and grown in full growth media unless otherwise noted for at least 24 h. Cells were washed with PBS and fixed in 4% paraformaldehyde for 15 min. Samples were then washed three times in PBS then incubated in blocking serum (1× PBS, 5% normal donkey serum, and 0.3% Triton X-100) for 1 h at room temperature. Primary antibodies were then diluted at 1:100 in dilution buffer (1× PBS, 1% BSA, and 0.3% Triton X-100) overnight at 4°C. Samples were then washed three times with PBS and incubated with fluorophore-conjugated secondary antibodies (1:100 in dilution buffer) for 1 h at room temperature. Cells were then washed three times in PBS and mounted onto glass slides. Cells were then imaged on an inverted Zeiss Axio Observer deconvolution microscope with a 63×oil immersion lens.

### Image analysis

#### Focal Adhesion Analysis

To analyze FA size and number, cells were fixed and co-stained with anti-paxillin antibody and phalloidin-Alexa596. For analysis, cells were randomly selected (using phalloidin-Alexa596 channel) in at least 5 image fields (typically 10-30 cells were analyzed for each experimental condition) using following criteria: (1) cell was not contacting any of the surrounding cells; (2) cell has clearly identifiable lamellipodia. All images were then acquired using the same exposure time. To select FAs, masks were created by image fragmentation and thresholding using anti-paxillin staining channels. The same thresholding and fragmentation criteria were used in all images. The size and number of the FAs was then measured using Intelligent Imaging Innovations 3I Imaging software. Data was derived from at least three independent experiments. Around 800-1000 FAs were analyzed for each experimental condition.

#### TAZ Activation Analysis

To analyze nuclear TAZ localization cells were fixed and co-stained with anti-TAZ antibody and phalloidin-Alexa596. For analysis, five random image fields (using phalloidin-Alexa596 channel) were selected, and all cells were analyzed in each field (typically 20-30 were cells analyzed for each experimental condition). All images were then acquired using the same exposure time. To select the nucleus, masks were created by image fragmentation and thresholding using DAPI staining channels. The same thresholding and fragmentation criteria were used in all images. The TAZ fluorescence sum intensity in the nucleus was then measured using Intelligent Imaging Innovations 3I Imaging software and expressed as fluorescence intensity per μm^2^. Data shown was derived from at least three independent experiments.

### Statistical analysis

Statistical analysis for all experiments was determined using GraphPad Prism Software (GraphPad). Datasets were assessed for normal distribution using the Shapiro-Wilk normality test. A two-tailed t-test was performed on all normally distributed datasets and a Mann-Whitney U test was performed for datasets not normally distributed. Data was collected from at least three independent experiments unless otherwise noted. In all cases, P ≤ 0.05 was regarded as significant. Error bars represent standard deviation unless otherwise noted. For all immunofluorescence experiments, at least five randomly chosen image fields per condition were used for data collection. For quantitative immunofluorescence analysis, the same exposure was used for all images in that experiment and quantified using Intelligent Imaging Innovations 3I Imaging software.

## Online supplemental material

Fig. S1 shows a sub-population of GFP-Rab40c colocalizes with Golgi.

Fig. S2 shows Rab40c antibody specificity, Western blotting and genotyping of Rab40c-KO cell lines.

Fig. S3 provides ANKRD28 mRNA expression levels in Rab40c KO cells, Western blotting of stable cell lines expressing FLAG-ANKRD28 or FLAG-PP6R1, and Western blotting of ANKRD28 knockdown cells.

Fig. S4 shows pFAK-s910 and the total FAK levels in Rab40c KO cells.

Fig. S5 shows all uncropped and unmodified western blots used in the figures.

Supplemental table lists all proteins identified in FLAG-Rab40c immunoprecipitation and proteomic analysis.

## ACKNOWLEDGEMENTS

We are grateful to Emily Duncan and Migle Prekeryte for critical reading and editing of the manuscript. This work was funded by the National Institutes of Health grant R01 GM122768 to R.P.

## SUPPLEMENTAL FIGURE LEGENDS

**Figure S1**

MDA-MB-231 cells stably expressing GFP-Rab40c were plated on collagen-coated coverslips and then stained with anti-GM130 (A; red) or anti-VAMP4 (B; red) antibodies. Boxes mark the region of interest shown in the inset. Arrow points to the lamellipodia.

**Figure S2**

(A) Testing specificity of anti-Rab40c antibody. 293T cells were transfected with empty vector, FLAG-Rab40a, FLAG-Rab40al, FLAG-Rab40b or FLAG-Rab40c. Cell lysates were then blotted with anti-FLAG or anti-Rab40c antibodies.

(B) Immunoblotting of cell lysates from control and Rab40c-KO cells with anti-Rab40c and anti-tubulin antibodies.

(C) Genotyping two of Rab40c-KO MDA-MB-231 cell lines. Deletions are highlight in red. Predicted amino acids are shown under the nucleotide sequences.

**Figure S3**

(A) qPCR analysis of ANKRD28 mRNA levels in control and Rab40c-KO MDA-MB-231 cells.

(B) WB analysis of lysates from MDA-MB-231 cells stably expressing FLAG-ANKRD28 or FLAG-PP6R with anti-FLAG antibody.

(C) WB analysis of lysates from MDA-MB-231 transfected with non-targeting control siRNA or siRNA targeting ANKRD28.

**Figure S4**

The effect of Rab40c KO on the levels of pFAK-S190. Cell lysates from control and Rab40c-KO cells were immunoblotted with anti-pFAK-S190, anti-FAK and anti-tubulin antibodies (A). Panel B shows quantification of the levels of pFAK-S190 phosphorylation. Data shown are the means and SEM derived from three independent experiments.

**Figure S5**

Uncropped and unmodified western blots used in the figures.

