## Supplementary figures and images for "Rab40c Regulates Focal Adhesions and Protein Phosphatase 6 Activity by Controlling ANKRD28 Ubiquitylation and Degradation"

### Supplemental Figure 1

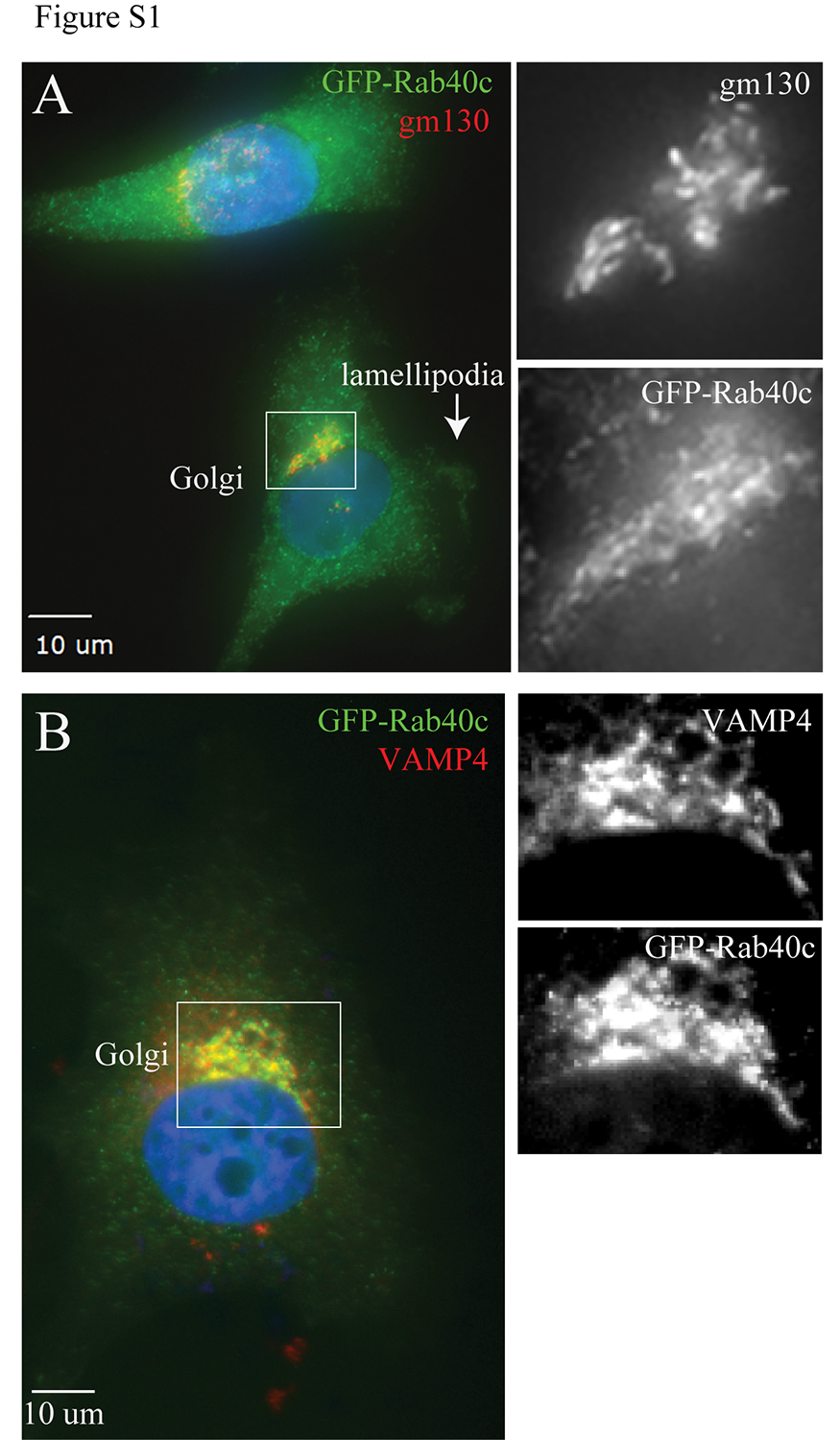

### Supplemental Figure 2

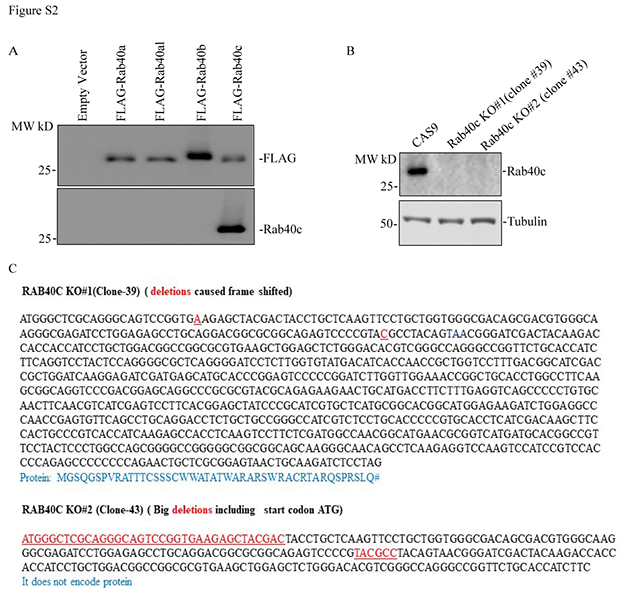

### Supplemental Figure 3

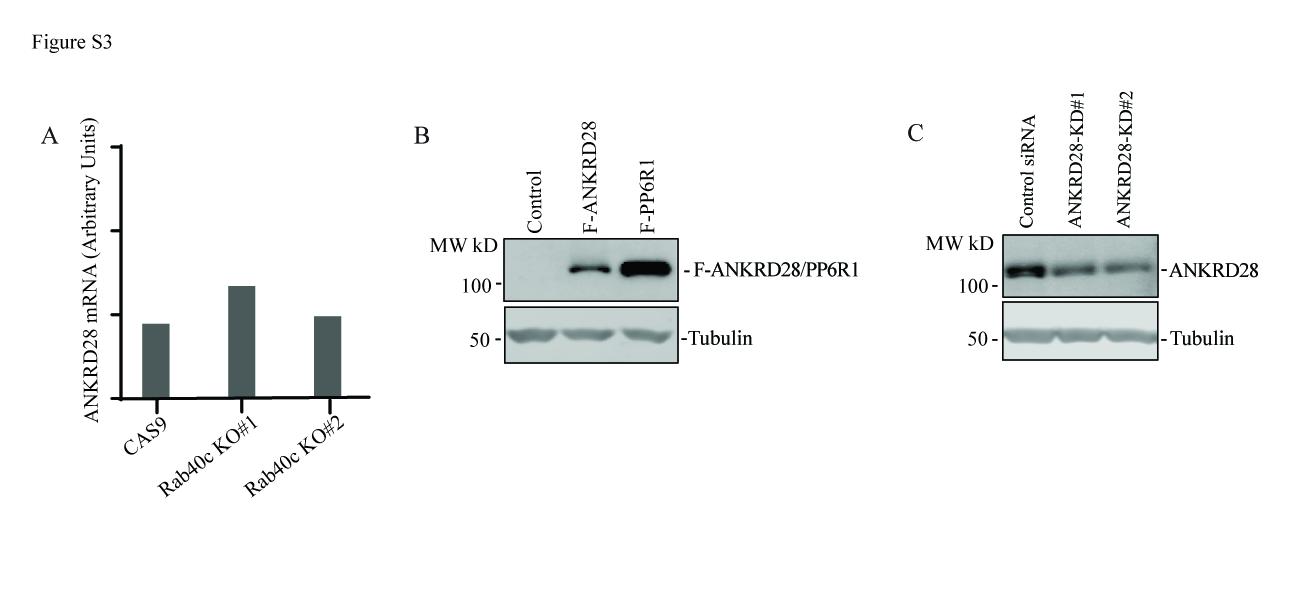

### Supplemental Figure 4

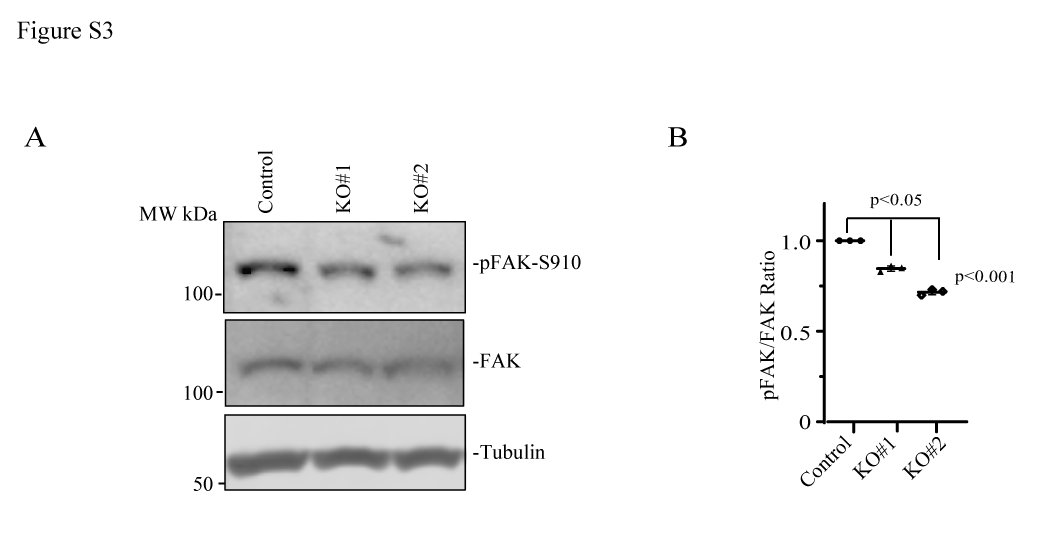
